# Software system design to support scale in mammalian cell line engineering

**DOI:** 10.1101/2025.09.01.673445

**Authors:** David W. McClymont, Baird McIlwraith, Althea Green, Sam Coulson, Elizabeth Scott

## Abstract

Cell line engineering (CLE) is the process of gene editing cell lines for a variety of purposes for research and development or bioproduction processes. Traditionally, CLE workflows have been manual and low throughput. Here, we describe the development of several software-based processes implemented alongside wet-lab automation and robotics, built to improve the throughput of our CLE platform to three times its previous capacity. A markup language (GEML) was developed to enable eCommerce capabilities and connections to internal manufacturing systems. A Laboratory Information Management System (LIMS) specifically designed to track CLE projects through all stages was created to manufacture the cell line specified by the GEML. Cell line engineering required analysis of images in brightfield without fluorescent staining therefore a machine learning (ML)-based method for analysing engineered clones imaged captured on an automated imaging platform was created. Our work demonstrates that combining both wet-lab automation and software approaches is essential to allow CLE workflows to reach their full potential, allowing the development of high-throughput robust platforms that meet the increasing demands of the field.

## 1. Introduction

Cell lines are defined as cultures of cells, derived from a single source, which can be propagated repeatedly in a laboratory setting, whilst retaining specific phenotypes and characteristics. Using various techniques, cell lines can be genetically edited to create new engineered cell lines, for example by knocking in or out of specific genes, to create disease models, or insert biological tags^1^. These engineered cell lines are important *in vitro* tools for R & D, drug screening, and bioproduction.

The discovery and subsequent development of CRIPSR/Cas9 gene editing technologies hailed a new era of precise, robust cell line engineering (CLE)^2,3^. Using CRISPR/Cas9 introduces double strand breaks (DSBs) in a cell’s DNA and can be used to create both knock out (KO) and knock in (KI) cell lines. Cas9 endonuclease is introduced into the cell, alongside a single guide RNA (sgRNA) containing a 20-nucleotide sequence which targets the Cas9 to the region of interest. After introduction of the DSB, the technology takes advantage of the cells own intracellular DNA repair mechanisms.

When generating KO cell lines, a process known as non-homologous end joining (NHEJ) is used. Repair of DSB by NHEJ is error prone, resulting in the introduction of non-homologous insertions or deletions (indels).^4^ These cause frameshift mutations in the DNA, preventing expression of the encoded protein. For KI cell lines, whereby a specific sequence is inserted into the genome, e.g. fluorescent tags, single nucleotide polymorphisms (SNPs) or whole gene sequences^5^, a donor DNA sequence is also introduced into the cell in order to make use of another of the cells internal repair mechanisms, known as homology directed repair (HDR) ^6^. The donor DNA sequence can be a plasmid, synthesised single stranded oligonucleotides (ssODN) or double stranded DNA (dsDNA). These DNA sequences contain overlapping sequences homologous of > 40bp to the site that the gRNA is targeting, known as homology arms, allowing the HDR machinery into incorporating the DNA sequence of interest into the genome.

To create an engineered cell line, therefore, there are a specific set of parts which can be defined. For editing using CRISPR/Cas9, these parts are as follows; a cell line e.g. HEK293, a guide RNA (sgRNA), a RNA molecule designed specifically to target the Cas9 protein to the gene or region of interest, a set or sets of PCR primers designed to amplify the appropriate regions for verification of the engineering event and, for knock in cell lines, a short DNA oligonucleotide sequence or plasmid.

Despite the development of CRISPR/Cas9 technologies for CLE, the practicalities of the physical workflow e.g. working with large number of cell lines at various stages of the process means that most CLE platforms remain low throughput. CLE pipelines include multiple complex stages, including transfection of the cell line with CRISPR/Cas9 components, single cell dilution (SCD) of transfected cells, clonal selection and genotypic validation of selected clones, all of which require tracking of constituent parts used, e.g. cell lines, sgRNA and primer sequences, and subsequently generated data, e.g. clone locations, images and sequencing data, through the use of databases or laboratory information management systems (LIMS).

Within synthetic biology, where design and engineering principles are paramount, many methodologies have been created to enable easy tracking of HT and large-scale processes and workflow, such as the establishment of specific language and definitions around engineering microbes. Concepts such as datasheets for reusable DNA ‘parts’ with standardised functionality have enabled the standardisation and reuse of parts in various genetic designs ^7^. The synthetic biology open language (SBOL) attempts to allow the communication of genetic designs in a graphical manner, and can be utilised using a graphical user interface (GUI) and connected to a LIMS for manufacture ^8^. SBOL also uses Glyphs refer to genetic circuit design and can map directly onto a specified sequence. DNA foundries have utilised the concept of designs and parts to define production of plasmids including verification of the end result ^9^.

Despite these learnings, very little has been done to attempt to translate this to mammalian CLE. Ontologies such as Cellosaurus have been developed to determine accession numbers for cell lines^10^ but, thus far, there are no standardised languages or definitions for the engineering of mammalian cell lines.

Here, we describe a set of cell line engineering platform innovations including of lab operations, automated data collection and analysis, data storage and development of standardised nomenclature, allowing for the implementation of an efficient, fully traceable, high throughput (HT) CLE platform. All research outlined in this paper is for research use only (RUO) not for use in diagnostic procedures.

## 2. Methods

### 2.1.1. Cell Line Engineering

CLE was performed via a standard workflow established within Revvity Discovery. Cell lines and CLE events were performed at the request of customers. Briefly, cell lines were transfected with Cas9, specific gRNAs and ssODN (if appliable). Transfected pools were then subject to limiting cell dilution in 384-well plates to produce clonal populations. Clonal populations in 384-well plates were imaged regularly to track clonality and growth rates. True clonal wells were then selected, a worklist created, and cells cherry picked using a Hamilton MicroLab STAR liquid handler (Hamilton). Selected clones were then genotyped via NGS to identify correctly edited clones.

### 2.1.2. Automated Image Acquisition and Analysis

To support HT CLE, an automated imaging workcell was designed, built, and integrated using Green Button Go (GBG) software (Biosero). The system consisted of a robot arm pf-400 (Brooks Automation), STX-520 automated incubator (Liconic), barcode reader (Keyence), plate hotels and a Cell Metric scanning imager (Advanced Instruments) (Figure 1).

**Figure 1:**
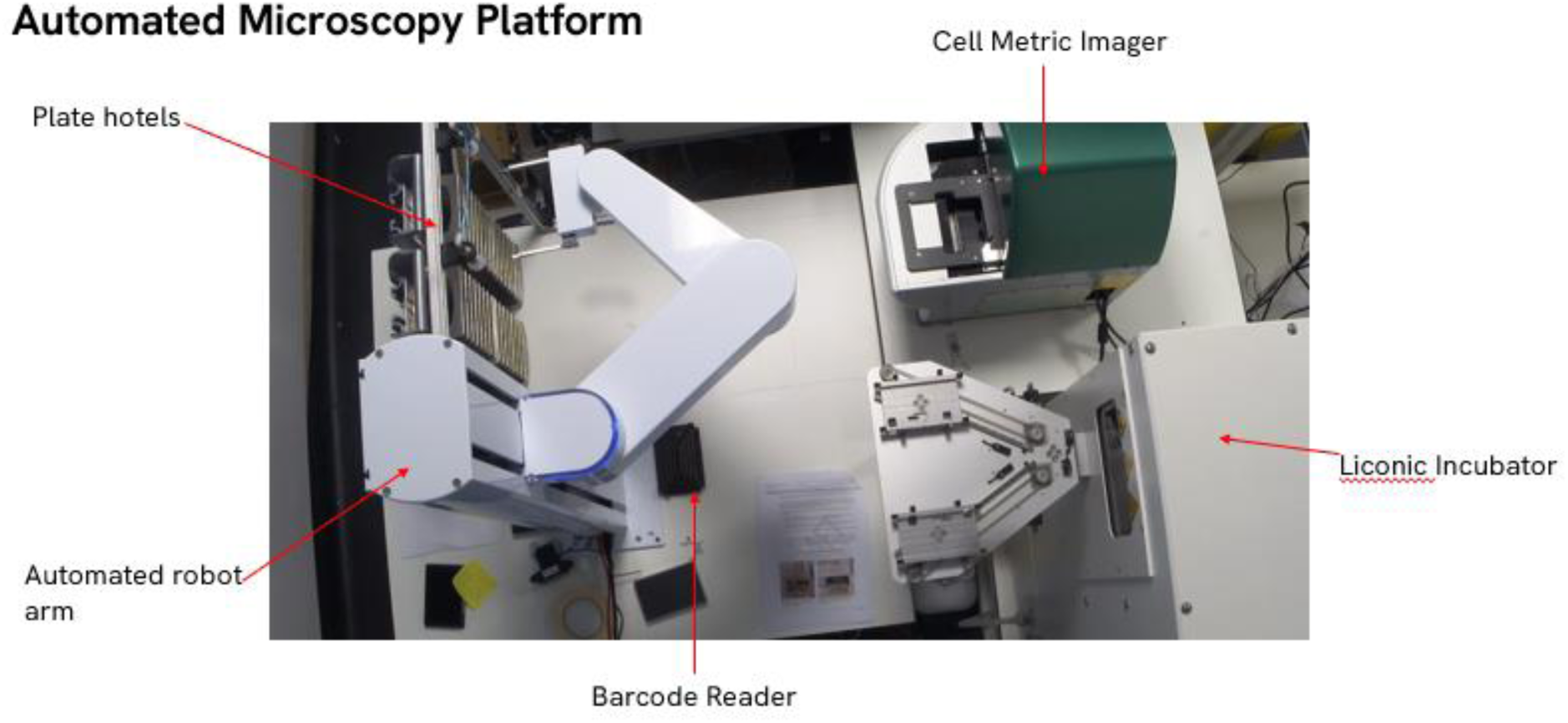
Automated integrated workcell for brightfield image capture system. The system comprises of a robotic arm which connects the automated incubator, barcode reader plate hotels and Cell Metric scanning imager for automated imaging of microtitre plates containing clonal populations generated from limiting cell dilution.

After limiting cell dilution, 384-well plates containing cells are registered into the system via the scanning of the individual plate barcodes and stored in the automated incubator. These plates were assigned to a ‘project’. The user submitted project numbers of be scanned to the GBG software at the start of a run. Briefly, the system then removed plates with barcodes contained within the project, one plate at a time, from the automated incubator and placed on the stage of the Cell Metric scanning imager for imaging. Once imaging is complete, plates were returned to the incubator. This continued until all plates from submitted projects were scanned.

GBG triggers a custom file uploader written in C# to upload images captured to the cloud. Ilastik^11^ was used to create the brightfield segmentation model. Images were then processed in the cloud using Ilastik headless mode and Azure Batch (Microsoft Inc.) for orchestration. The data processing pipeline automatically scales virtual machines based on need. Resulting data was stored in Cosmos DB (Microsoft) where a web application was written using asp.net framework for interaction with the data.

### 2.1.3. LIMS System design and implementation

For the design and implementation of a LIMS system for the HT CLE platform, a set of minimum requirements were drafted, including availability of CRISPR design tools, Electronic Lab Notebook (ELN), inventory, sample management system and dashboard building capability. Request for proposal were sent to vendors, resulting in the selection of Benchling (Benchling Inc.).

Data models for cell line, reagents and the CLE process were created and mapped into Benchling ‘Entities’. ELN templates were created for various stages of CLE (Transfection, SCD and Expansion) and pages were linked together into a process on Benchling using a request/execution cycle. Dashboards for tracking metrics were written in SQL and ran on the Benchling Insights package. Our CLE service website and ordering service was connected to the Benchling LIMS and its output converted to GEML using azure function apps and azure service bus messaging service.

## 3. Results

### 3.1.1. Data Model Development

A data model was developed for cell lines allowing CLE to take place with full tracking of each cell at any stage, whilst generating a standard ontology for identification of cell lines and the engineering outputs at various stages of the process. The starting point for cell lines used for engineering are frozen batches with the lowest possible passage, as long term culture can cause undesired alterations in the genome, metabolism or protein expression ^12^. Therefore, the ability to track and deconvolute those properties is desirable particularly for unexpected or undesired genetic changes or contaminations. Implementation of this data model allows the development of a LIMS to then manage and track that manufacturing process to completion.

### 3.1.2. Cell Line Data Model

Cell line entities were linked in a 4-level hierarchy to ensure consistency in naming conventions and to maintain correct connections to their parental cell lines (**Error! Reference source not found.**2). The highest level was the canonical cell line object, which contained the cell line name from Cellosaurus and was matched 1:1 with accession numbers from the Cellosaurus database. Cell lines that are genetically the same or similar have a wide variety of common names that make standardisation and data management difficult. For example, the cell line ‘HCT 116’ can be referred to as HCT-116, HCT.116, HCT_116, HCT116, HCT116wt, HCT-116/P, HCT-116/parental and CoCL2. The Cellosaurus ID was instead used to group together all common names under one accession number. All cell line names which were contained within a single accession number in the LIMS under the metadata field ‘Aliases’. The data model correctly grouped together cell lines in a way that is robust, scalable and searchable within the data model for all common names used. Users were unable to enter this as free text in the LIMS system, ensuring standardisation.

The second level was ‘Cell Bank’, which represents a particular source or clone of a given cell line. The entity has a parent-child relationship with a Cellosaurus ID and obtains its name from the name field of the parent object. The cell bank level enabled two sources of the same cell line to be represented. For instance, HCT 116 from provider A and provider B may be genetically identical but as they came from a different source with no way to determine the provenance therefore must be represented within the data model rather than the two cell lines being considered the same.

The third level, ‘Cell Bank Batch’, represents the batch and lot number of a given frozen vial on receipt at our site. It contains metadata related to physical processing, such as date banked and passage number. Cell bank batch allows issues related to a particular vial to be tracked if there were issues with freezing or contamination related to the specific vial used.

The fourth level, ‘Cell Chassis’, represents a physical batch of live cells in culture. A batch of cells in culture can either be used for CLE or propagated for further use, with an associated increase in passage number. Each ‘Cell Chassis’ entity created at this level represents a culture of cells used to make an engineered cell line, and this incremented as required if growing cells were taken at different points. The term chassis is commonly used in synthetic biology as the host of any engineering parts in microbes ^13^. We transposed this term to mammalian cells growing in culture and gave an ID number based on passage.

The structured identification code therefore follows the pattern of Cell bank ID, Cell Bank Batch ID, and finally the Cell Chassis ID of the cells (Figure 2). For example, CBB045-002-001, represents the first chassis and second batch of cell bank 45. The combination therefore represents all information required to pinpoint the exact provenance of the cells used in a single CLE project. The cell line data model was developed to capture and deconvolute the provenance of any cell object in the system considering their ability to constantly expand.

**Figure 2:**
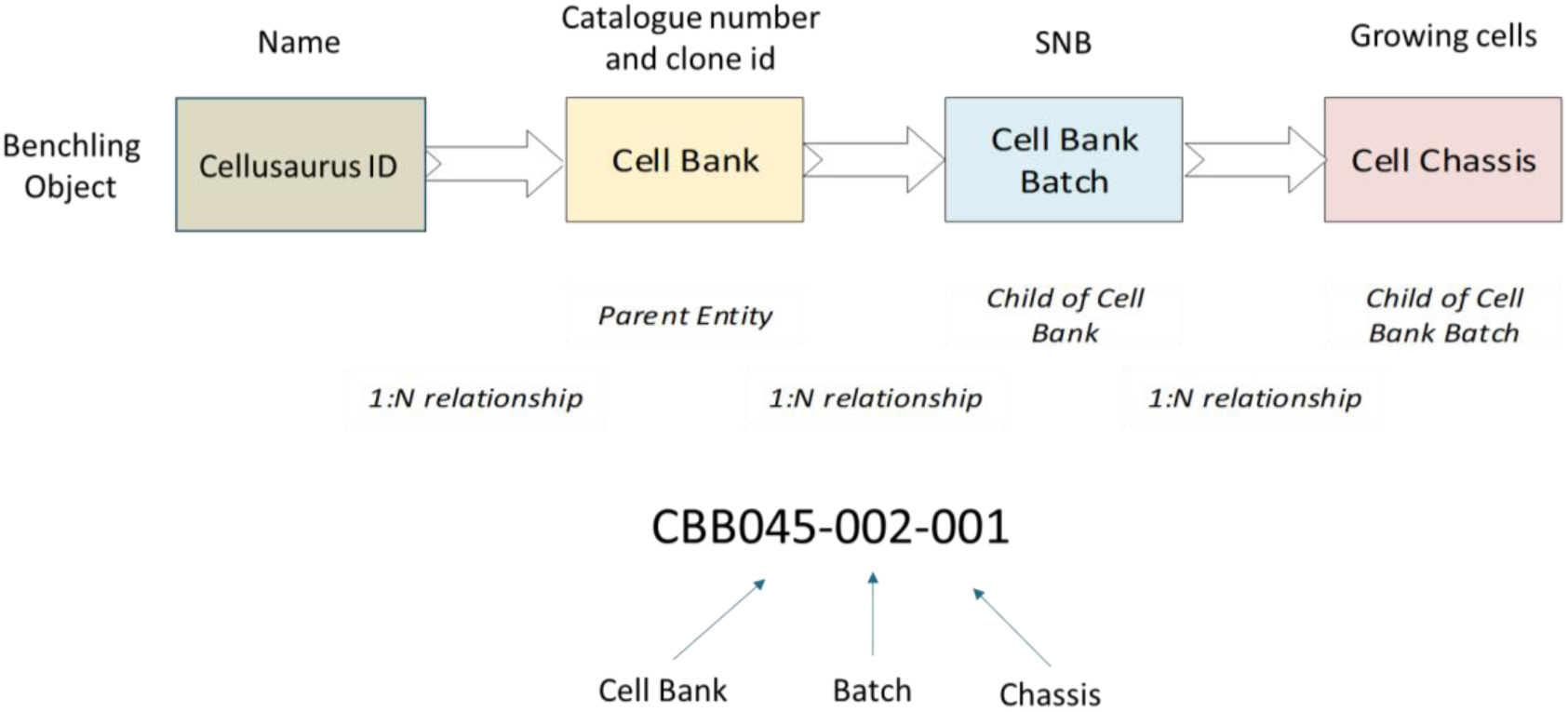
Entity relationship diagram for the newly generated Cell Line Data Model. Each cell line is assigned a unique ID using a 4-level hierarchy, including Cellosaurus ID, Cell Bank, Cell Bank Batch and Cell Chassis.

### 3.1.3. Reagents Data Model

Reagents used in CLE projects, such as Cas9, sgRNAs, ssODNs, plasmids and primers, followed a simple hierarchy and they were given a parent and child batch level connection. The parent entity represents the immutable characteristics of the object, such a DNA sequence or species, whereas the child batch level represents the specific instance of that object, such as lot number, source or concentration. The standardised naming system developed for these entities followed a typical naming convention, using the ID of the parent object with the number of the child batch appended to it e.g. gRNA001-05.

Specific forward and reverse primers were designed and created for each CLE project and used to sequence the area of the genome has been targeted during the CRIPSR/Cas9 editing event. Individual primers follow the parent-child batch method described for reagents. A primer pair meta-entity brings the forward and reverse primer into one grouping. The primer pair id is then linked to the gRNA ID, representing the specific area that will be amplified (Figure 3).

**Figure 3:**
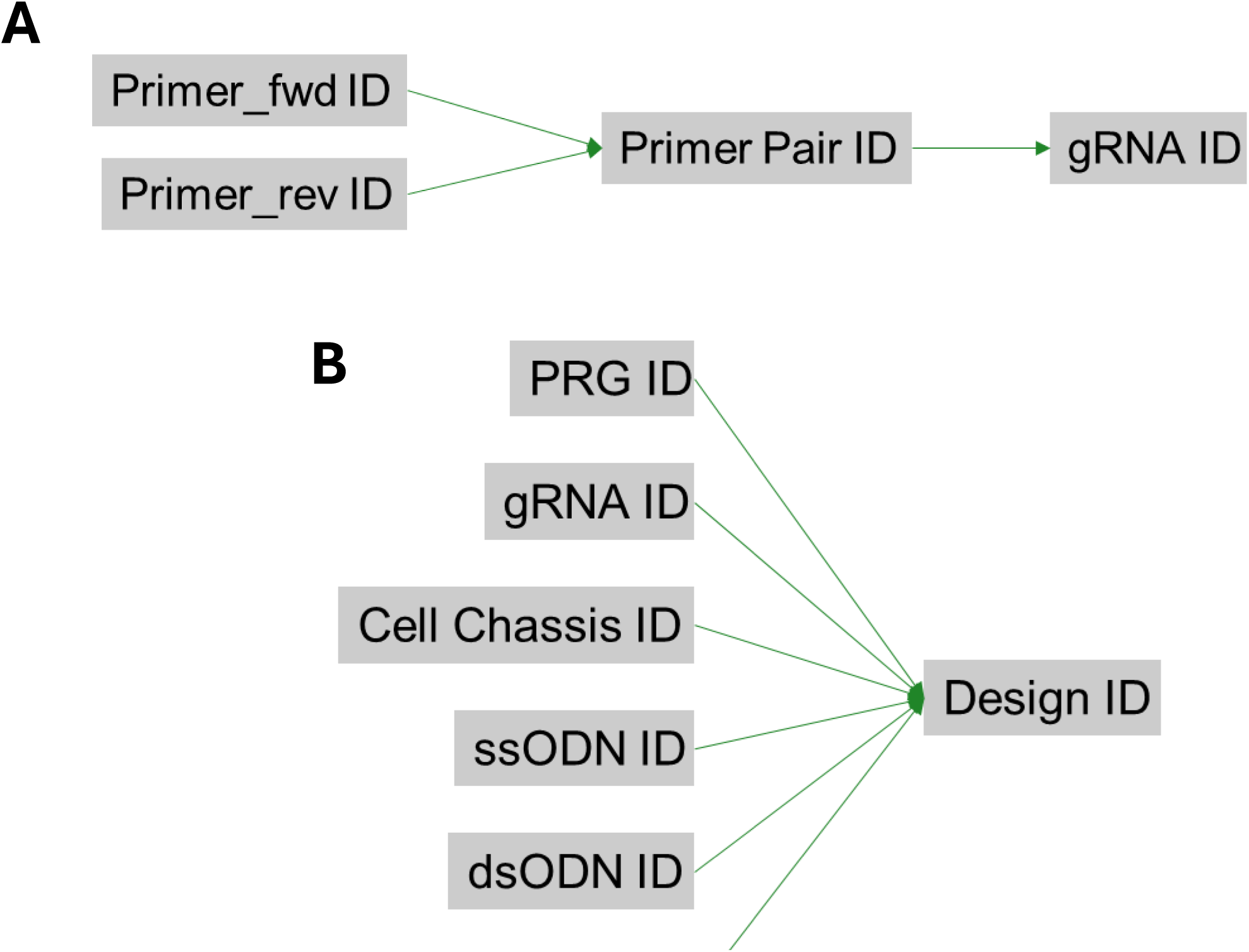
Grouping meta-entities within the data model. (A) Entity relationship diagram for primers and primer pairs data model. A forward and reverse primer is brought together as a primer pair id that is linked to the gRNA for which the region is amplified. (B) Entity relationship diagram for design data model. The design ID brings together specific instances of all the entities required to build a cell line engineering project. PRG ID refers to a program or project ID that relates to a specific cell line deliverable. gRNA ID is the guide RNA targeted to a specific genomic location and Cell Chassis ID is an instance of cells in culture that have been taken for engineering. The remaining DNA parts enable knock-in projects encoded with homology and the modification desired. DNA parts are single stranded oligonucleotides (ssODN ID), double stranded oligonucleotides (dsODN ID) and plasmids (Plasmid ID).

**Figure 4:**
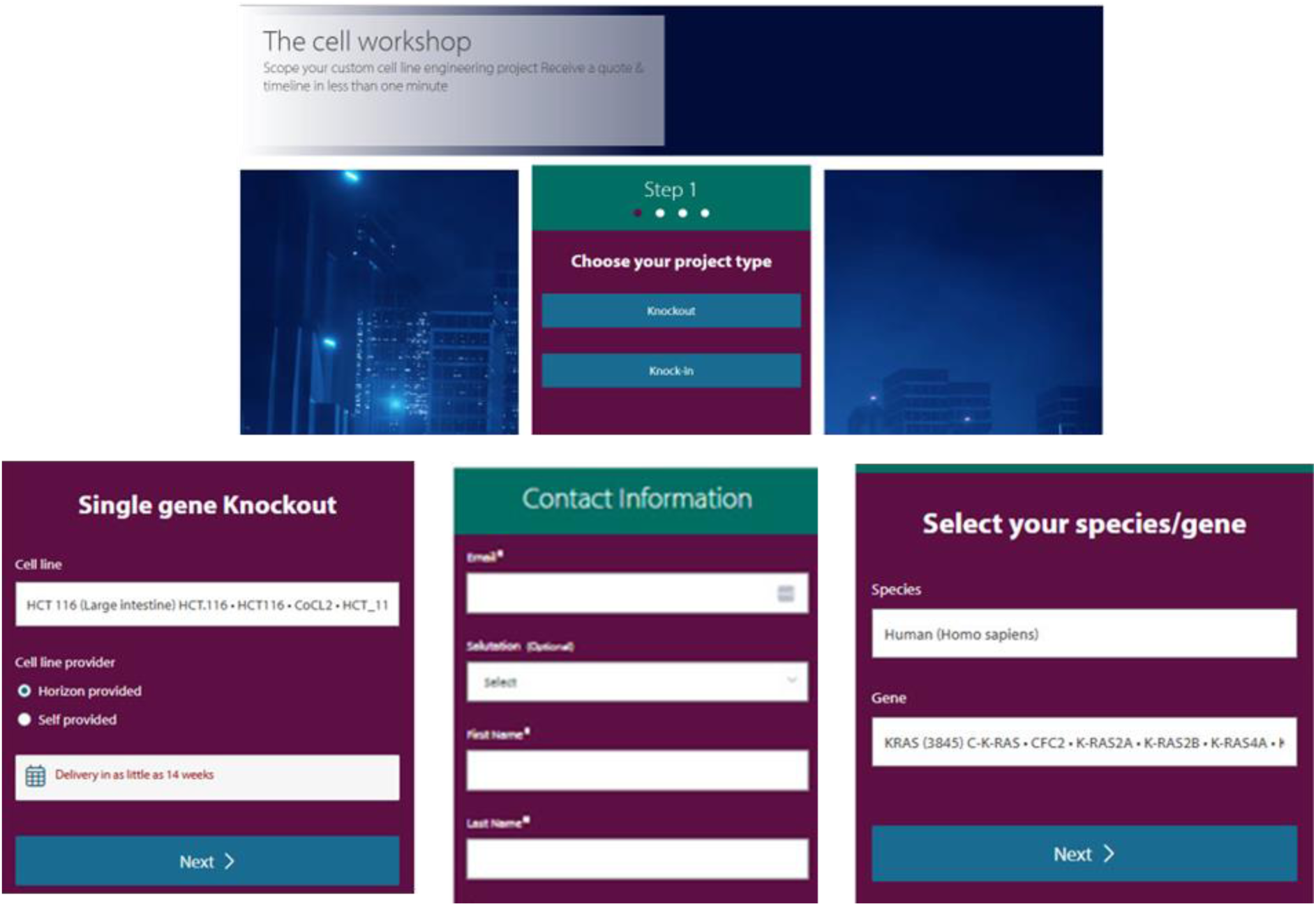
Screenshots from CLE Configurator tool. Customers entered data at each stage detailing the specifications for their desired cell line.

### 3.1.4. Design Data Model

To represent the combination of cell lines and reagents within the LIMS, a grouping entity known ‘Design ID’ was created to bring those entities together into a single meta-entity (**Error! Reference source not found.**3). A programme ID (PRG ID) refers to a cell line deliverable for a given customer and contains connections to IDs in other systems, such as eCommerce.

### 3.1.5. Creation of Genetic Engineering Markup Language

To have a robust data model that encapsulates the complete set of entities required for a single CLE project and ensures cell lines are created to the correct specification, would require manually creating entities, as well as the correct connections between entities. Registering a typical project would need creation of 60 entities on average, with 139 entity links. Manual entry of this information directly would a single user >1 hour.

A method was therefore developed to allow end users to configure their gene edited cell line on the website themselves and automate the process of order entry and entity registration into the LIMS. A structured markup language was created to ensure robust immutability in the messages being passed between systems and to ensure that projects were registered correctly to produce the desired CLE result.

An XML format called Genetic Engineering Markup Language (GEML) was created that combines the sequences of the sgRNA and primers, and cell line with the correct hierarchical connections to register a design. GEML is a canonical, immutable, unambiguous design specification for how to build a cell line from a series of parts. The XML is enclosed in GEML tags which encloses all subsequent tags used for the project; <Order>, <Engineering> and <Manufacturing>. The <order> tag encloses information such as customer name and sales order ID. The <engineering> tag takes all the elements related to the CLE process such as engineering type (KO or KI) and sgRNA and their related primer sequences, as well as metadata associated with those sequences. The final <manufacturing> tag section outlines information relating to the final product, which is produced, including cell line information and various deliverables such as number of clones to ship to the end user.

When an order was created on the website using the configurator to define the requirements (**Error! Reference source not found.**4), the end user enters species, cell line and the gene target, as well as their details for delivery. Doing so resulted in the creation of an Azure Service Bus message that records those details, and this is then processed into a sales order. The sales order is then processed by an Azure function app and converted into GEML. A service bus message containing the GEML was picked up by a subsequent function app. This app registered all the required entities in LIMS with the appropriate connections in place, circumventing the need for time consuming and error-prone manual data entry.

### 3.1.6. Implementation of an efficient LIMS allows for efficient and robust reagent management

After developing an efficient way of registering all entities associated with a CLE request, we then created a system to accurately manage and track the CLE process itself. Our CLE platform is divided into multiple phases: Initiation, Electroporation, SCD and Expansion. Each phase creates new, related entities, depending on the physical process being performed. Using the LIMS feature ‘Requests’, entities associated with a specific ‘Request’ are placed in a workflow within a ELN template and executed by processing though the tasks in the ELN page.

For example, to initiate a CLE workflow, a workflow request was made for a cell line to undergo electroporation, this request contained links to all associated design entities, and the ELN page tracked the processing of those entities. After electroporation, linked entities called ‘Pools’ were created, which have a parent entity of the design ID. These ‘Pools’ are an entity which represents a mixed population of cells with a variety of editing events. ‘Pool’ entities that are considered successful via sequence verification, i.e. identification of the correct target sequence within the pool, are then included in a request for SCD. Seeding of selected pool entities are then executed and tracked through a new SCD ELN page. At the end of process, another set of linked entities called ‘Clones’ are created, with a parent relationship to the ‘Pool’ they were derived from. The ‘Clones’ are again verified by sequencing and those identified as having the correct target sequence, are included as entities in a request for expansion. The expansion phase consists of growing cell cultures in larger vessels to produce cell numbers sufficient for banking in cryovials for shipping to the end user, and this is also tracked through the use a specific ELN page.

### 3.1.7. Development of a Request Execution Cycle for HT CLE project management

The request execution process described above, accommodated quality control (QC) verification, rework loops and dead ends at any stage of the process, and enabled high levels of flexibility to run many possible pathways simultaneously. A single CLE project can have multiple designs, producing many child ‘Pool’ entities. Only successful ‘Pools’, i.e. those containing cells which have been correctly edited, move to the next stage of the process, with the unsuccessful pools being discarded. The same occurs for ‘Clones’ derived from those ‘Pools’ - many ‘Clones’ are created from a single ‘Pool’ however only those with the required gene editing event are expanded to be shipped as product.

After assessment of the ‘Pools’, new designs can be created and executed if no ‘Pools’ are identified as successful. For example, new designs could have the same sgRNA with a different cell chassis, or completely new sgRNA with the same cell chassis. Multiple ‘Pools’ can be sent via request to the SCD, or could be performed multiple times with the same ‘Pool’ if there is enough material. ‘Pools’ can also be used as cell chassis’ themselves on new designs, either with the same sgRNA to increase editing efficiency, or with a different sgRNA for additional editing at a different gene locus. A ‘Clone’ can also be used as a cell chassis for a subsequent CLE project, accommodating multiple gene edits and tracking all stages of the process *n* levels deep. All lineages and rework routes are tracked within the same program due to the hierarchical natures of the connections between the entities and their execution through the requests system (Table 1).

**Table 1:** Hierarchy and linkage for entities at each stage of the process.

| Level | Type | Execution Stage | Parent |
| --- | --- | --- | --- |
| 1 | Program | Initiation |  |
| 2 | Design | Electroporation | 1 |
| 3 | Pool | SCD | 2 |
| 4 | Clone | Cell Expansion | 3 |

### 3.1.8. Robust CLE project monitoring using LIMS dashboards allows for accurate and efficient workflow tracking

To allow scientists and management to have an easy and efficient way to view the status of individual CLE projects, we developed a series of Dashboards using the Benchling Insights subsystem. Dashboards are backed by the SQL editor, which we used to pull all the relevant data pertaining to a project from across the LIMS platform. Results, entities and ELN page statuses were queried, bringing all the different strands of a project together into one concise view. The dashboards also provided direct clickable links to the relevant entity or ELN page, allowing users to easily view the details of a specific CLE project if required.

Dashboards were created for each phase of the CLE process (Initiation, Electroporation, SCD Cell Expansion) (Figure 5). Each dashboard was made of multiple blocks; one for ‘Incoming Requests’, showing a list of those requests waiting for action, one block containing ‘Request Execution Info’, detailing the ‘in progress’ ELN pages with status, a block which showed ‘Outgoing Requests’, a list of requests from the perspective of the current stage to the next, and the final block ‘Re-Executions’ showed requests considered to be repeats or rework loops. Each row in the block links to other pertinent information associated with those requests, such as due dates and project name.

**Figure 5:**
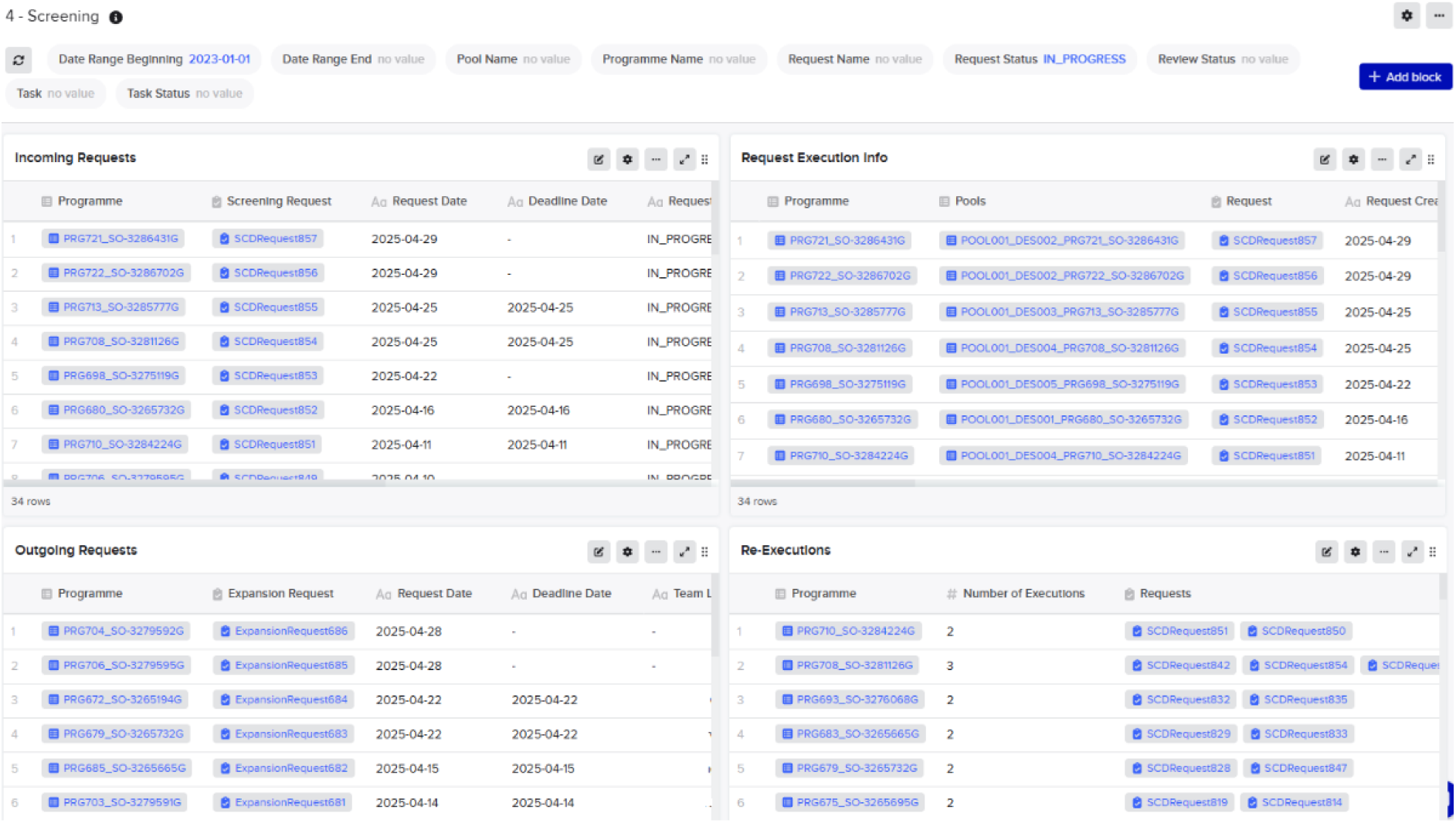
Examples of Dashboards created for managing Single Cell dilution and Cloning stage of the process. The four quadrants help manage the process of incoming requests (top left), executed requests linking to the associated ELN page managing the process (top right), outgoing requests represented clones selected to move to the next stage (bottom left) and re-executions which highlights those executions that are from repeated projects (bottom left). A range of filtering options include data ranges and execution state enable focussing the dashboard to the desired information

Additionally, a Dashboard was also created to show platform level metrics for all active projects, known as the Project Overview dashboard. ‘Days Since Last Stage’ was a measure used to accurately and reliably track how many days had elapsed since a project had entered the most recent phase in its request execution cycle (i.e. Initiation, Electroporation, SCD or Expansion). Projects that were stuck at a specific stage of the process for an prolonged period could then be easily identified. To get this information, the number of days that had passed between the current date and the creation date of the relevant request was obtained using SQL. ‘Furthest Stage Reached’ defined the furthest stage in the request execution cycle that the project had reached in the overall workflow. Due to rework loops, the ‘Furthest Stage Reached’ was not always the stage the project was currently in. Therefore, we defined an additional metric ‘Latest Stage Reached’ to allow users to easily see which phase of the CLE process a project was in.

The graphing capabilities of Benchling Insights Dashboards were also used to provide high level visual metrics for management of our HT CLE platform (Figure 6). Metrics included project completions per week/month, average repeat rate per project and request execution stage, average project lead time, number of repeats per project type and average time spent in stage. The programme metric dashboards were SQL queries connecting data from across the Benchling ecosystem, such as request dates, entity metadata fields and Results table contents. Repeats were calculated by counting number of requests in each stage of the request execution cycle. The time spent in each stage was returned by calculating the time delta between the creation and completion of a specific ‘Request’. Data needed to be binned by month, which was achieved by extracting the day and month of each data point, then grouping and sorting the data by those dates. There were also date filters for the periods of interest and project types. Overall, development of multiple Dashboards using the Benchling Insights Dashboards feature, allows for us to build reliable tools for accurate tracking of high numbers of CLE projects within out HT CLE platform.

**Figure 6:**
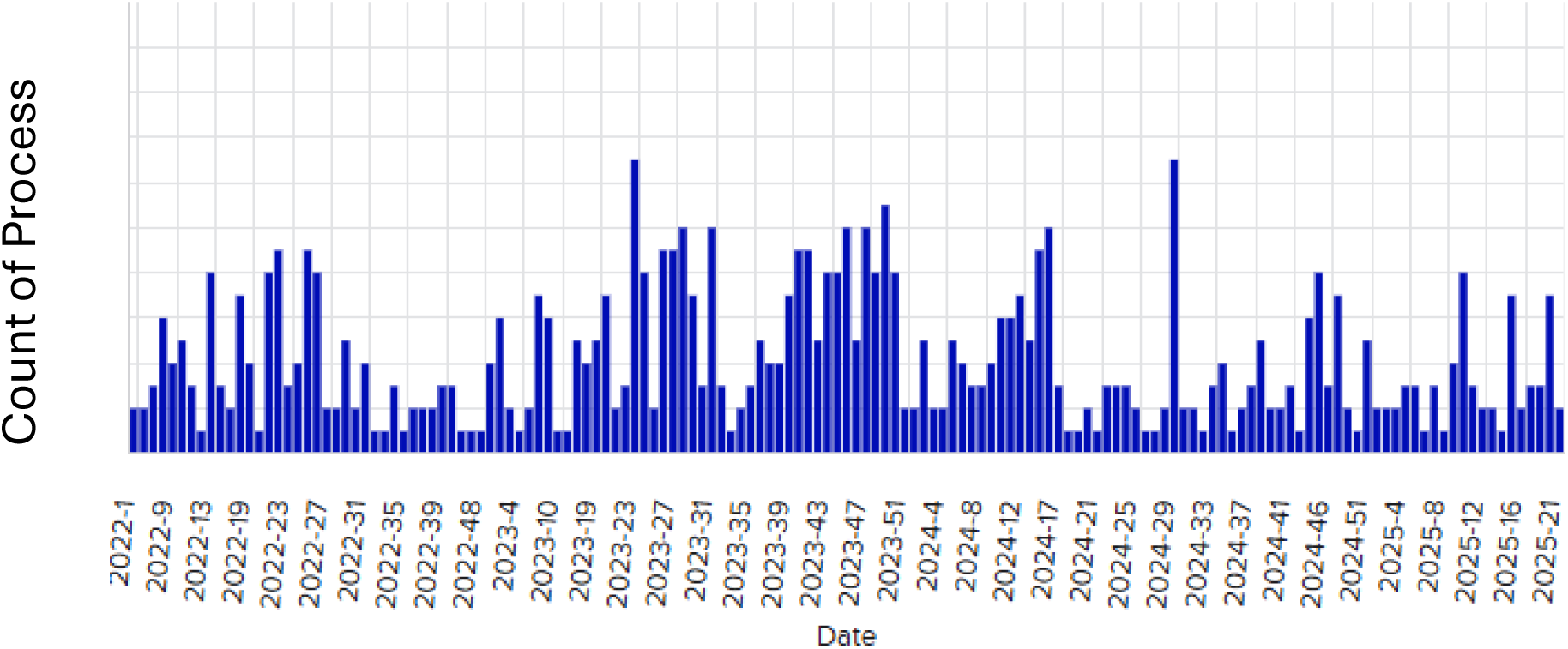
Dashboard displaying an overview of number of processes carried out per week over time

### 3.1.9. Reproducible and automated generation of CLE datapacks

On completion of each CLE project, a datapack was generated for delivery to the end user. Datapacks are PDF documents containing details about the Cell line generated during the CLE project, such as parental cell line information, sgRNA and primer sequences used for CLE, and sequencing data. The datapack allowed the customer to see all QC steps performed to ensure the edited cell line meets the manufacturing specification, and that the CLE process has been performed correctly, as well as details about other reagents that can be ordered for further experiments.

As the LIMS system contained a data model that tracked the entire lineage of all entities during each phase of the CLE process, information could be gathered by working back from the final clone. The clone is the end node of a tree and all information required for a datapack from one piece of information from the lowest point in that tree. To do this, a Benchling template was created, and the Benchling Lookup function was used (Figure 7). In a Lookup Table, the leftmost column acts as a ‘key’ that can then be used to link to any related entities in the entire hierarchy. A ‘Clone ID’ can be used to look up the ‘Pool’ and ‘Design ID’ entity it was created from. Once the Design ID entity is found, the linked guide sequence and Cell line information can be located.

**Figure 7:**
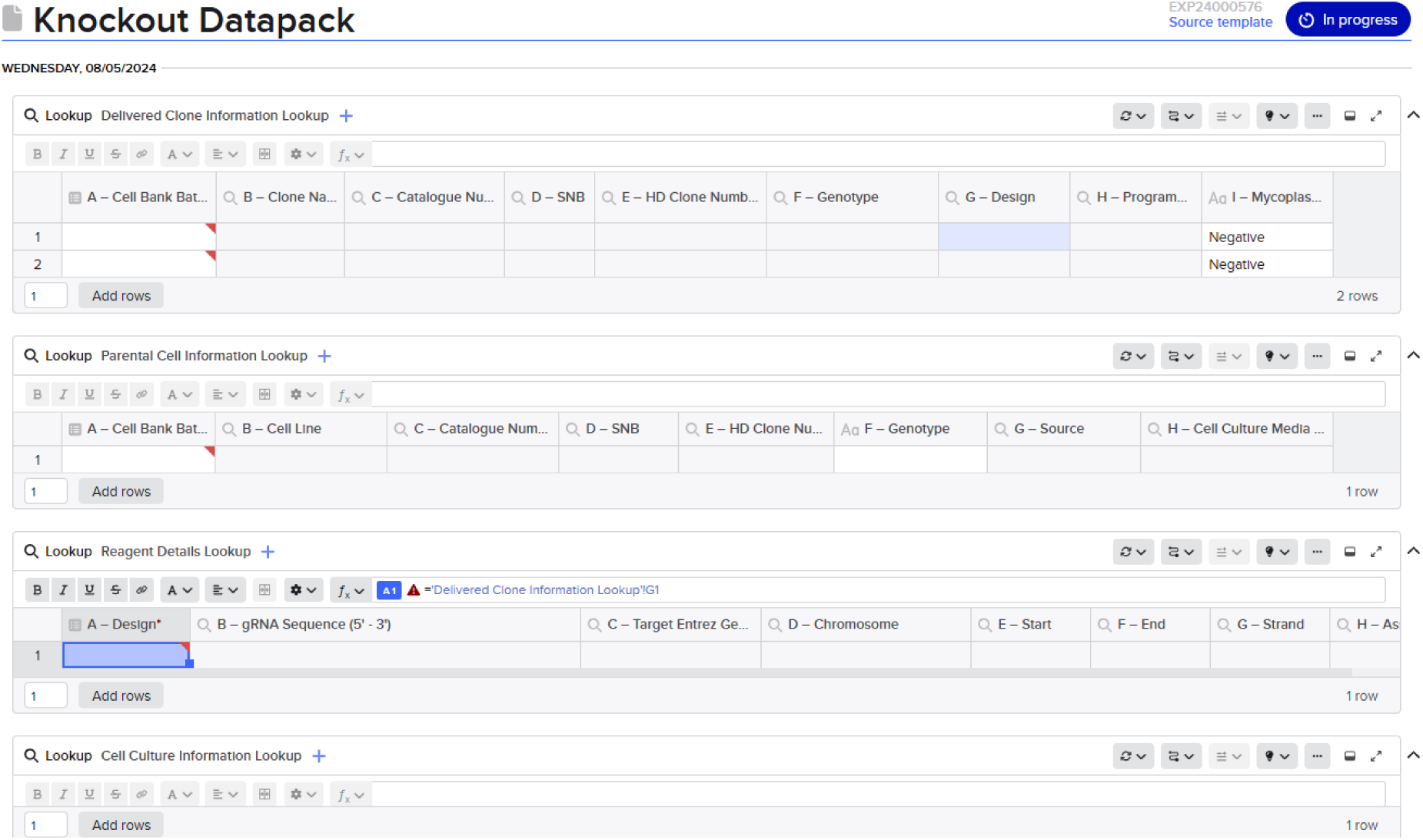
Datapack template layout. Entry of a cell bank batch which represents a completed cell line engineering project displays linked metadata in the tables that is required for the datapack. Subsequent tables then use the metadata for further lookups, chaining them together and enabling all data to be collated from one cell bank batch ID. Datapacks are then exported as a PDF to send to the customer.

The Lookup Tables were placed in an ELN page and multiple lookups were chained together into multiple tables, displaying all the data required in a single template page. By chaining tables together, all data required for the datapack was brought together onto page with the user only having to enter one entity. The ELN datapack page was formatted to be customer facing containing corporate logos and was exported as a PDF and sent directly to the customer. Production of a datapack by a scientist or manager took approximately 2 hours and, therefore, development of this automated process for producing CLE datapacks significantly increased efficiency, whilst reducing the likelihood of errors.

### 3.1.10. Custom Software Development

While the LIMS was developed by configuring a commercial software platform, for processes that are not universal, custom software needed to be built. One of the largest bottlenecks in our CLE process was within the SCD phase of the process. After limiting cell dilution of cells in 384-well plates, brightfield images of each well are taken at regular intervals to track clonality. The bottleneck comes from assessment of these images to produce a list of wells with clonal populations, including excluding empty wells, and those derived from multiple cells.

Development of our automated integrated imaging platform helped alleviate the bottleneck in collection of the images. Often, immunostaining is used to aid segmentation in microscopy, aiding the highlighting of cells as the background of the well is not stained and therefore black. However, fluorescent stains can be toxic, therefore, brightfield techniques are preferred in CLE processes ^14^. Using traditional segmentation methods on brightfield images leads to high noise and variability, making it difficult to analyse wells with sufficient resolution ^15^. Therefore, we developed ML image analysis model and a custom web application to remove this bottleneck.

### 3.1.11. Ilastik ML Model for accurate identification of clonal populations

We developed ML image analysis models using Ilastik to improve the signal to noise ratio and segmentation capability in the brightfield images of individual wells. These were then orchestrated into a cloud-based application for processing and interrogation. The robustness of the model was tested to ensure its suitability and benefits over existing methods. We had previously found that main source of noise within segmentation-based imaging methods was false positive edge detection. Within the greyscale brightfield image the change in intensity at the edge of single well within a 384-well cell culture plate was classified as cells with other, more basic analysis methods (Figure 8). When tested on an empty 384-well cell culture plate, however, the Ilastik model measured no confluence on almost all wells tested.

**Figure 8:**
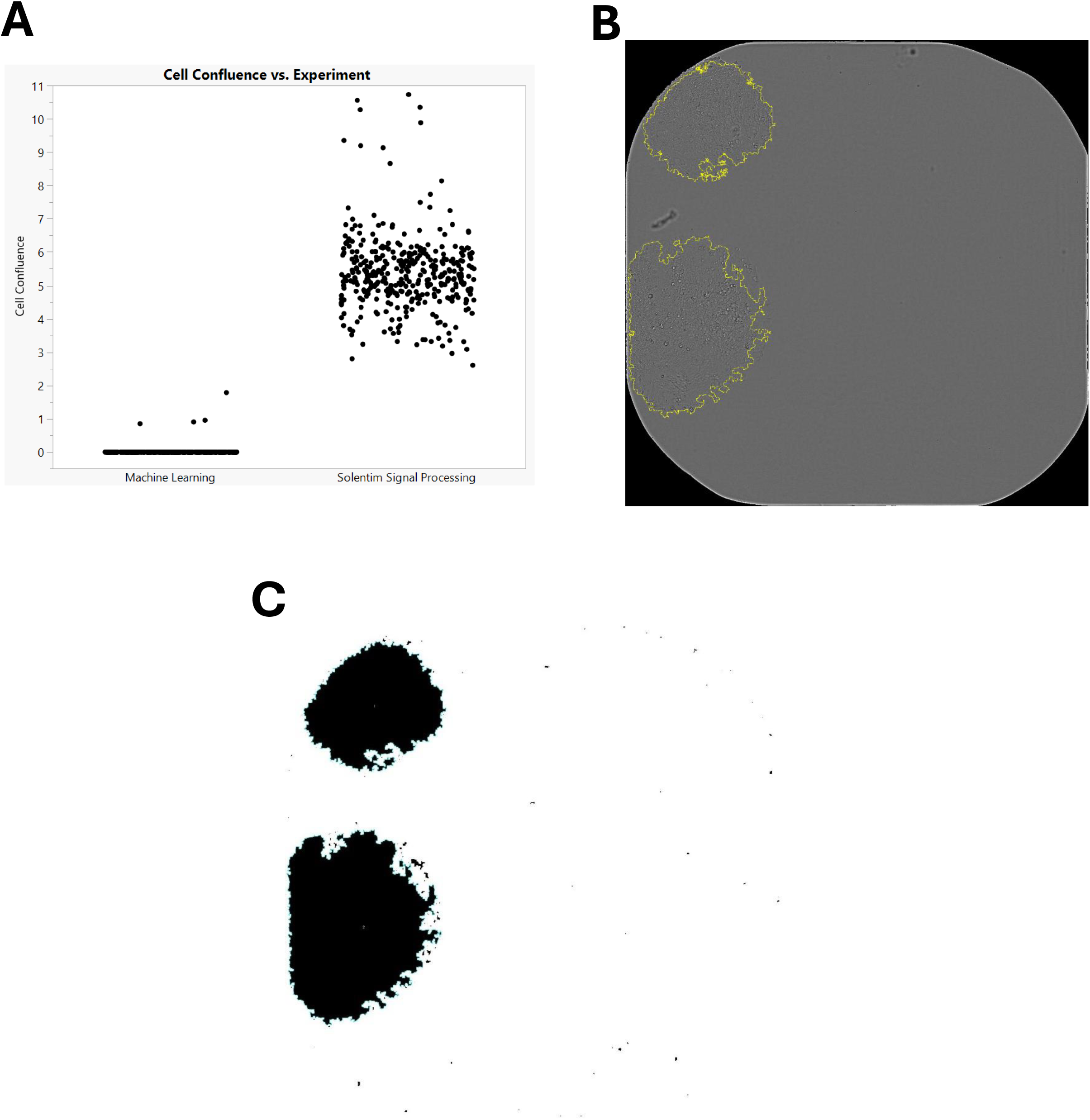
(A) Percentage confluence measured in empty 384-well plates measured by the capture instrument (Solentim Signal Processing) and our Ilastik ML model. (B) Example image of one well of 384 well plate with object detection overlay. C) Classified imaged by the Ilastik model used for object detection and colony counts.

To test the sensitivity of the system for detecting low cell numbers in these wells, the lower limit of detection was determined. We demonstrated that small colonies i.e. those identified as being less than 1% confluent could be detected across multiple cell lines (**Error! Reference source not found.**2). In classified images, areas containing cells were identified, with limited-to-no detection of cells in areas of the well where there were no cells present, even at low confluence. Other possible scenarios were also explored to test the lower level of detection and robustness with cell doublets or single cells that land in proximity within a well (Figure 9). Due to the nature of limiting dilution, doublets and single cells that are close to each other could be misinterpreted as clones, particularly if the read frequency was not high enough (Figure 9). Cell lines including HAP-1, HCT-116 and HEK-293 were tested over time, with tests showing that the analysis model was able to successfully detect the increasing size of colonies over time (Figure 10).

**Figure 9:**
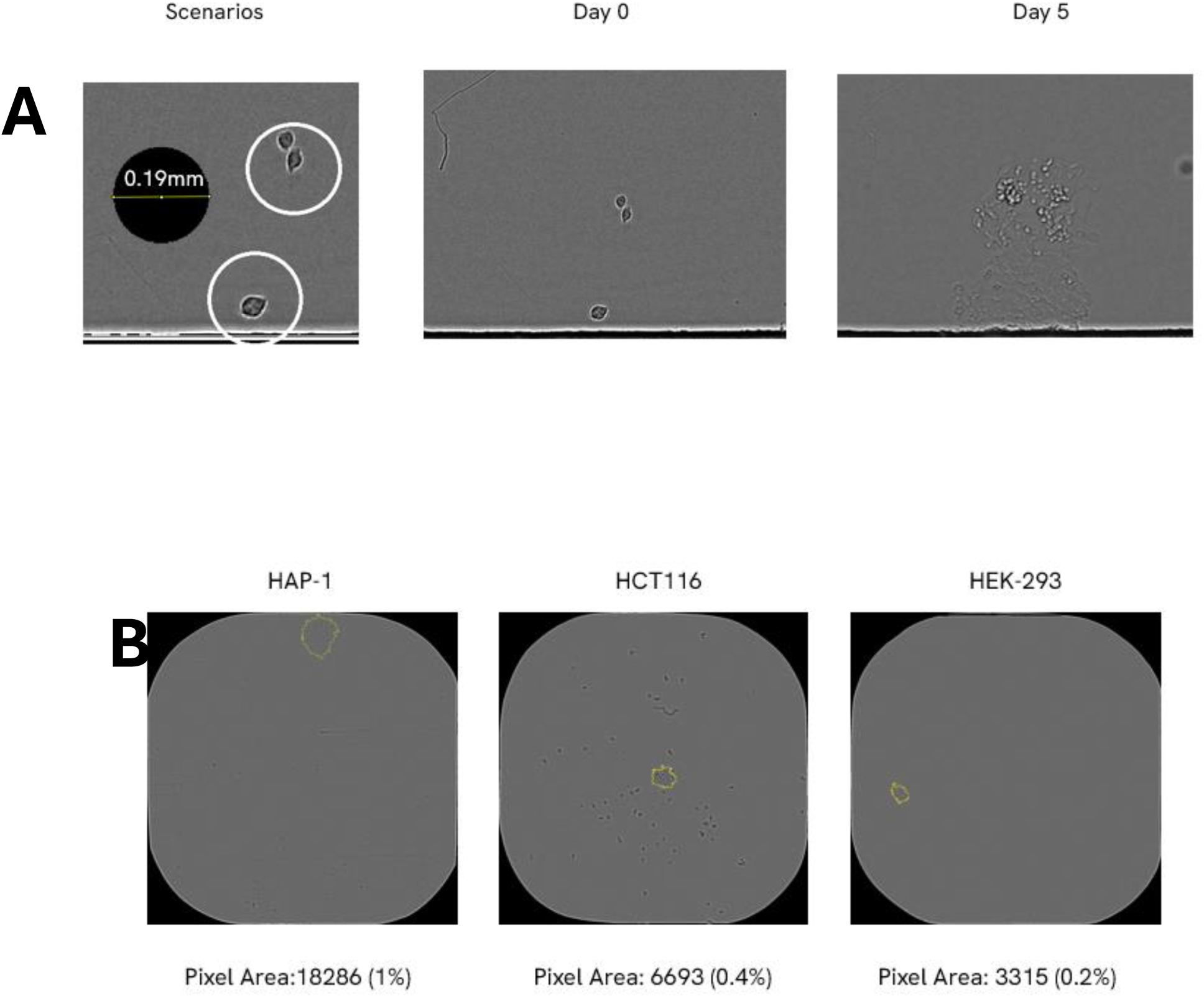
Example Images and Scenarios. (A) Doublets and cell proximity Images. Due to the nature of limiting dilution, doublets and single cells that are close to each other could be misinterpreted as clones, particularly if the read frequency was not high enough. (B) Lower limit of detection across cell lines. Small colonies with less than 1% confluence could be detected across multiple cell lines and non-cell material in wells not detected as colonies.

**Figure 10:**
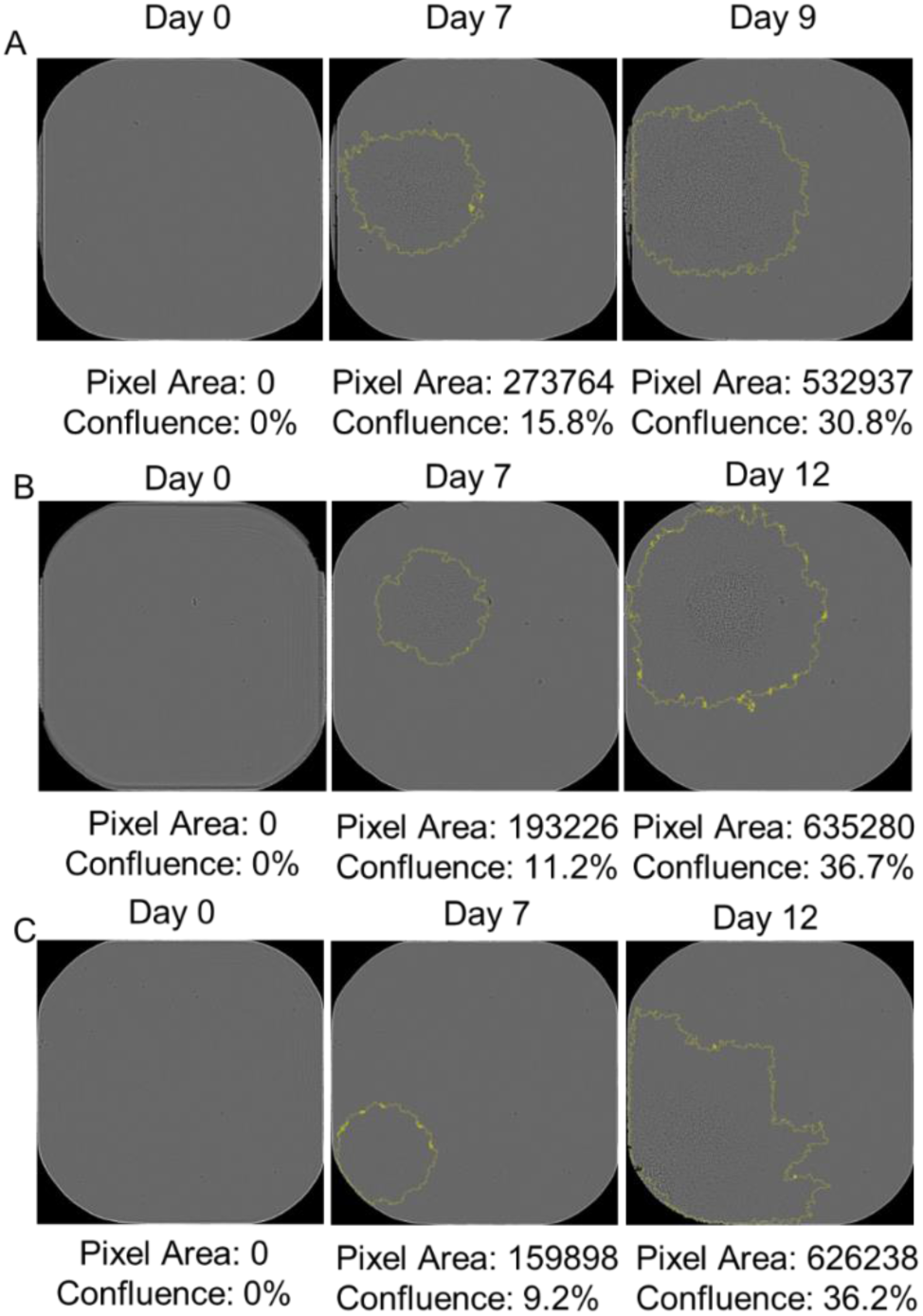
Example images showing detected colonies across time course experiment. A shows Hap1, B shows HCT116 and C shows HEK-293

**Figure 11:**
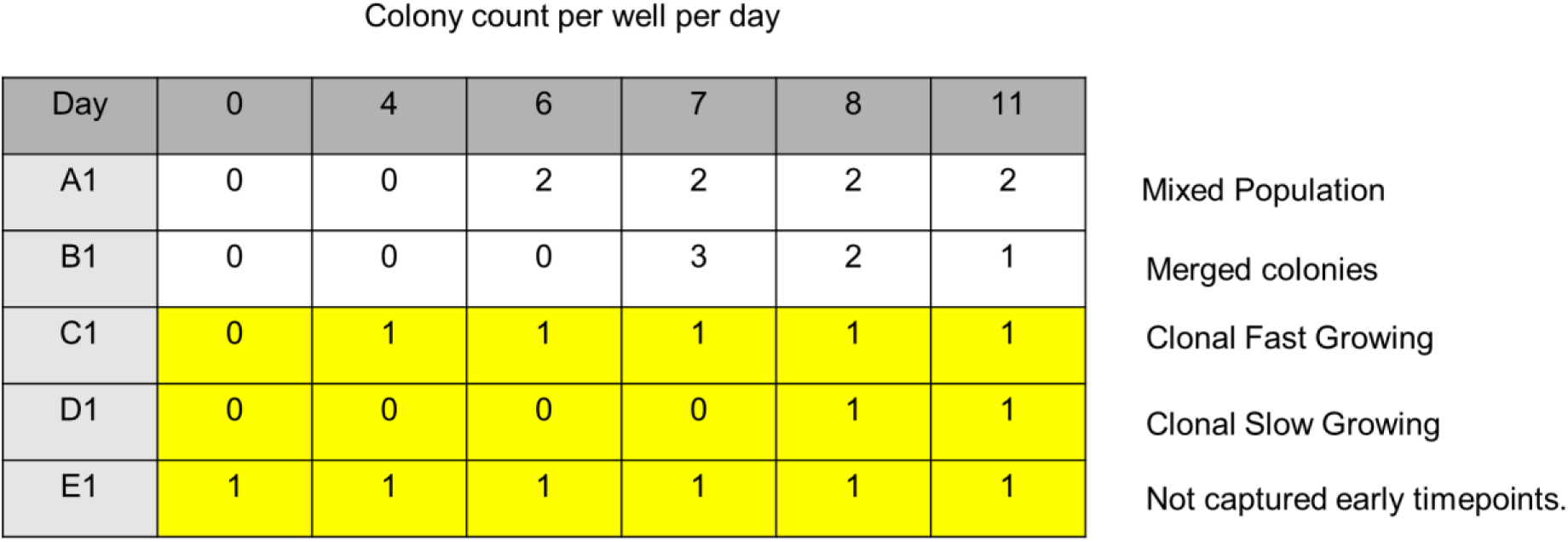
Truth table outlining the colony counts per day that are considered a clonal population. The state changes are tabulated and the desired state changes that represent a clonal population of cells is highlighted in yellow.

**Figure 12:**
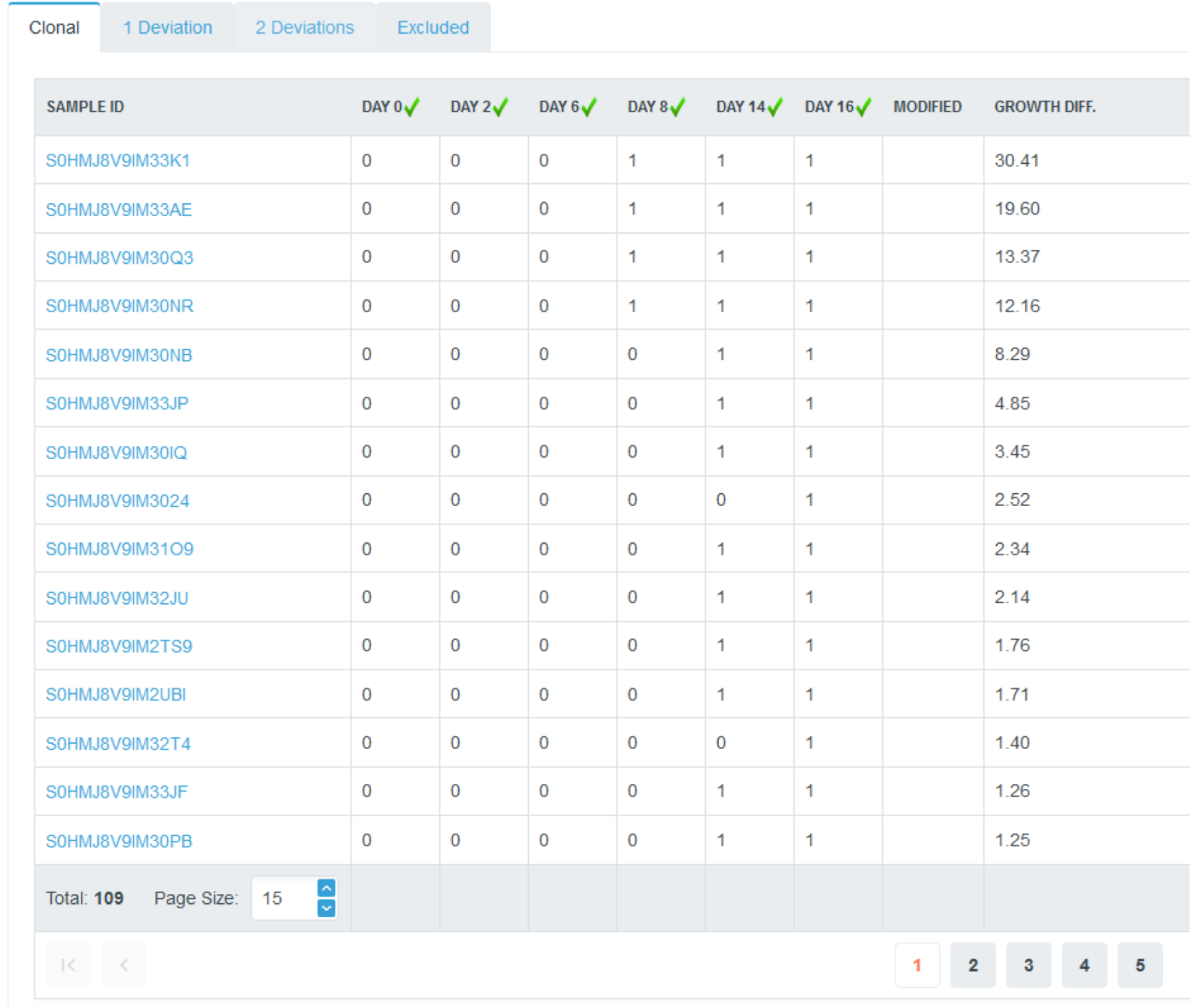
Study view showing list of samples and the colony counts over time. Samples for the entire pool are split into tabs representing different state change over the timecourse measured. The tab selected is the clonal population with allowable pairwise states 0-0 0-1 and 1-1. The list is sorted by colony size. Tabs for 1 and 2 deviations from the desired states can be selected and manually QC’d.

**Figure 13:**
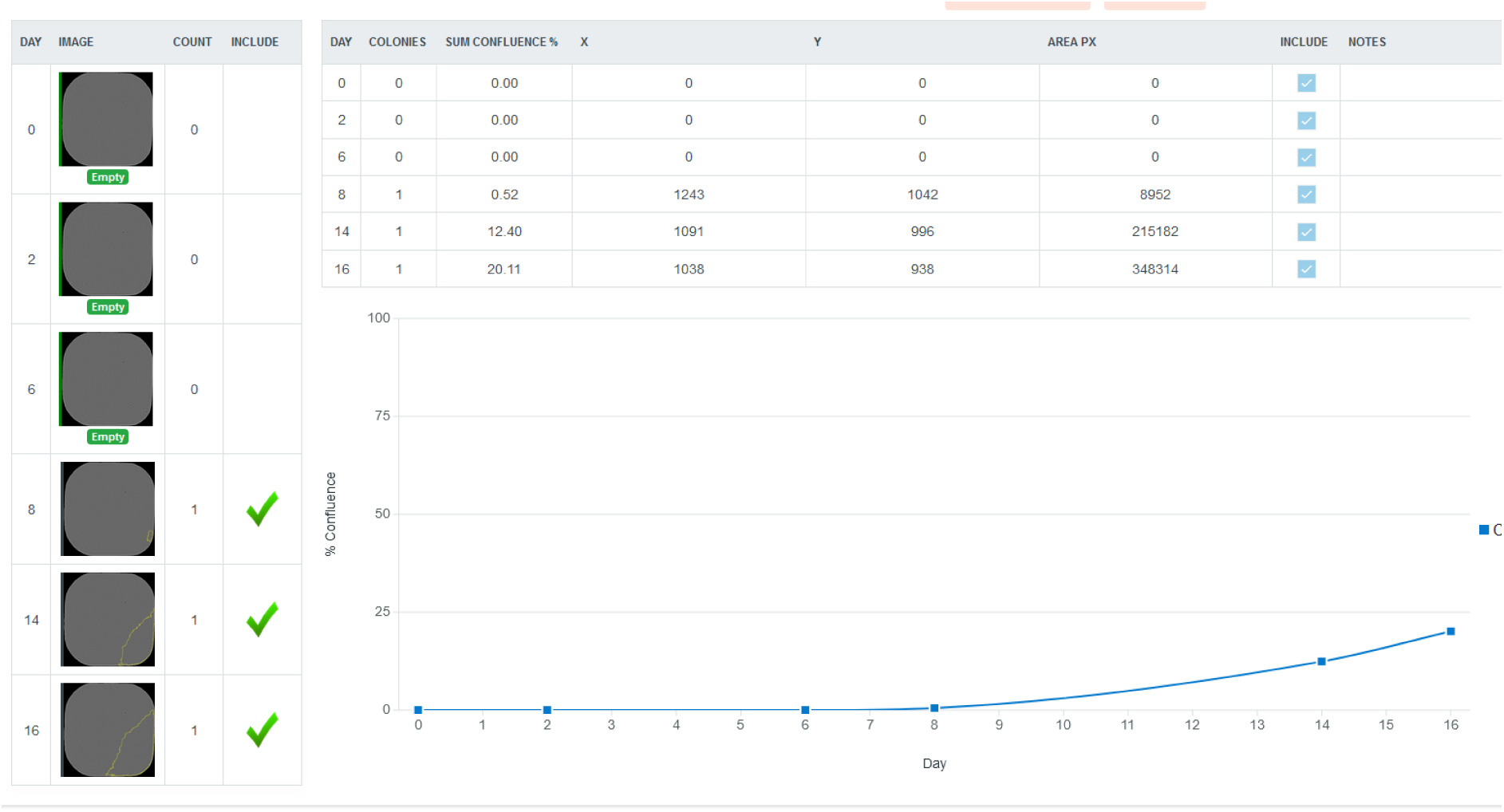
Sample view shows images from a specific well captured over the entire time-course. The Table shows individual colonies which can be included or excluded from analysis. Line connecting points show confluence increasing over time.

For each image of the wells taken, the system created a classified image using the Ilastik model that detected cells and not well background. From that, a black and white was generated contiguous shapes that represented colonies could be detected. The count of discrete closed shapes then represented a colony count, and this was recorded per time point at which well imges were collected. Colony count is the mechanism used to exclude wells and define wells as clonal and therefore wells with wells containing >1 colony were excluded from further analysis. As colonies grow radially from a single cell, with a low enough sensitivity, deriving colony counts can then be used quickly to exclude those wells, ensuring the final worklist contains only wells containing single colonies or ‘clonal’ populations.

### 3.1.12. GBG Automation Workcell Cloud Integration allows for efficient automated data transfer

For a given CLE project, on average ten 384-well plates were created within the SCD phase of the workflow, each with their own unique barcode. The capacity was 740 plates, therefore it was essential to develop methods to group plates together to trigger the capture of images of the necessary plates on specific days.

We first developed a method using the GBG software that registered groups of plates within the system. Plates that contained cells from a specific ‘Pool ID’ which had been subject to SCD were placed on the hotel. GBG then prompted the user to enter pool ID and the robot arm moved plates from the hotel to the barcode reader to be scanned. The barcodes were then written to a .csv file named according to the [Pool_ID].csv entered by the user. The rows of the .csv file contained all plate barcodes associated with the pool id.

A second GBG method takes a list of Pool IDs to scan plates required a given day. A .csv file was produced containing a list of Pool IDs to be scanned. For each Pool ID, it searches the folder that contains all the [Pool_ID].csv files available in the system. If one is found that matches, then the barcodes contained in that file are enqueued into a picklist file using the GBG function ‘Enqueue Barcode.’ The barcode list allowed all barcodes from all Pool IDs to scanned in the microscope using the robot arm in the running process.

A custom .dll library was created to enable the interaction between GBG and Azure. The .dll was placed within the installation folder of GBG allowing C# scripts executed by a GBG process to include and run Azure functions. After a plate has been read on the microscope, the read plate event function is executed that writes a service bus message containing the Pool ID and barcode of the plate. A file uploader service running on the microscope capture PC subscribes to that message and uploads the image to Azure File Storage. The Uploader service then sends a service bus message registering that upload that is subscribed to by an azure function app that process and enqueues the plates in the Cobra Workflow function app. Using Azure Batch SDK, it orchestrates the creation of parallel VMs running the Ilastik analysis. Batch nodes are configurable to process n plates in parallel representing one VM per plate.

### 3.1.13. Cobra Web Application for automated generation of worklists

When images of wells were analysed using the Ilastik model analysis pipeline, a list was produced containing confluency and colony counts per well. A methodology was then developed to obtain a list of wells that the pipeline had determined were ‘clonal’, i.e. only derived from a single cell, which could then be used downstream by an automated liquid handling system for cherry picking. Colony counts were tabulated under several scenarios to determine what states changes over time would be considered ‘clonal’ (**Error! Reference source not found.**1). By considering colony counts pairwise, shifting by one position across, state changes 0-1 and 1-1 with leading 0-0 are allowed; whereas, all other state changes result in exclusion from the clonal set. Therefore, by using a simple rule set, all other scenarios representing undesirable outcomes are excluded e.g. multiple cells in wells, colonies that merge and are mistakenly counted as one colony or colonies which are lost over time.

A software platform was developed that allowed scientists to produce a worklist of clonal wells, and the ability to interrogate images and QC data, known as COBRA. The main interface view showed the user a list of studies that were currently active in the system, with a study defined as a set of cell culture plates (>10) which contain cells that have undergone SCD. A table shows the date the set of plates were last imaged and the difference between the largest and smallest confluence to give an overview of those studies that need to be actioned.

Focusing on an individual study, a list of all wells was shown with the colony counts over time. Samples that only have state 0-1 or 1-1 are listed under a tab called ‘Clonal’ (**Error! Reference source not found.**2). The sample view was able to exclude, in bulk, all images captured on a specific day if there were issues processing images or problems with the optics on the instrument. Further tabs also showed lists of samples where clonality is undetermined i.e. they may be clonal or may show unallowed pairwise state changes. These are termed deviations, and there are tabs listing samples with one or two pairwise state changes outside the allowable set. The ability to recover samples near expected value allows QC and interrogation to recover samples that may have been false negatives in error.

Individual samples could also be interrogated in further detail in a per-sample view. This view brought together all the images captured from a single well over time. The output from data processing at each stage is tabulated colony size/confluence of each colony at each timepoint. Individual colonies can be excluded or included which can make the overall output more accurate if required (**Error! Reference source not found.**3). The clonal state is automatically calculated, so if any exclusion or inclusion changes the overall state in such a way that its grouping changes, it will be added to the appropriate list.

After QC, a list of sample ids that are considered clonal is used to produce a pick list that can be downloaded in CSV format and used for further processing by an automated liquid handler. The sample IDs are converted into the physical barcode of the 384-well cell culture plate and the well which they are located int. Studies are then marked as complete and cannot be altered further. Across 429 studies 80% were above the cut off value of 96.

## 4. Discussion

The development of CRISPR/Cas9 technologies for mammalian CLE has revolutionised gene engineering. However, despite this, CLE platforms are often low throughput, relying heavily on manual tracking and data input. We have developed CLE eCommerce and platform innovations to allow the implementation of high-throughput gene engineering workflows orderable *via*. website. A fundamental data model was created that unambiguously defined what a CLE project was so that it could be codified into XML that enabled communication between various software systems. The data model also enabled the creation of a comprehensive LIMS to manage and track the manufacturing process. eCommerce platforms for biotechnology applications have been limited to basic catalogue items or simple unambiguous data models e.g. DNA synthesis or cloud labs. The work presented here describes a unique manufacturing on demand (MOD) system managing large numbers of stock keeping units (SKUs) of all possible genomic engineering events in cell lines, enabled by existing and future technologies.

Within the website, customers specified engineering event attributes such as cell line and edit type. These attributes were captured via the CLE Configurator Tool, which aimed to replicate the simplicity of the customer facing web-based interfaces used to connect the eCommerce systems and automated labs producing synthetic nucleotides. For synthetic nucleotides, capturing sequences is simple as customers often know exactly what sequence they require, so therefore it can be easily captured on an online form and transferred to a manufacturing system queue^21^. GEML built on this concept and extended it to the more complex process CLE. Standardisation enabled robust connections to other systems such as enterprise resource planning (ERP) and LIMS, initiating all required workflows necessary to sell and manufacture an engineered cell line. The GEML defined a specification for the outcome of the design that any sequencing results could be measured against.

Synthetic biology combines engineering with biology to focus on forward engineering of desired properties using designs and parts to achieve that aim. Typically focussed on single celled organisms with established and mature technologies, mammalian synthetic biology has lagged in similar standardisation despite the promise of more human relevant modalities such as CAR-T and theranostic cells ^16^. Therefore, the approach taken when developing the CLE platform was based on applying synthetic biology principles to our process particularly parts, designs and data models and applying them to mammalian synthetic biology^17^. For editing using CRISPR/Cas9 technology, the target loci within the genome are encoded in the specific sgRNA sequence. Therefore, a design that collects the nucleotide reagents required (sgRNA, DNA and primer pairs for subsequent sequence validation) designed for the specific targeting event required come together with the Cas9 enzyme and cell line as parts. GEML codifies the parts together with the engineering blueprint that is mapped onto the data model for manufacture. Doing so gives a repeatable, scalable language and infrastructure brining the principles of synthetic biology to mammalian CRISPR cell engineering.

LIMS systems are an important tool to help scientists manage and schedule work efficiently while recording data required for decision making and process improvements. A centralised database creates a single source of truth where emergent useful features are possible, resulting in new ways of working and realising increased capacity compared with disparate excel spreadsheets. As much as possible, data entry through the processes was designed to be passive rather than active, so data within the models was created by running of the process itself. Doing so within a centralised database enabled features such as automated generation of data packs for customers as all entities and data were linked together in the model. The hierarchical connections of the data model meant that the entire tree of information associated with a cell line could be obtained from a single piece of information immediately.

The request-execution cycle of the LIMS system represented the stage gates where child entities are created i.e. those where the cells undergo a transformation or selection of some kind such as transfection or expansion. At each stage of the process, the requests could be executed again, allowing for tracked re-work loops seamlessly built into the model and created by carrying out the project within the system. The ELN pages used to manage the process at each stage were templatised. However, multiple templates or ad-hoc changes to that page were allowed to build in flexibility to the process e.g. different types of transfection or data analysis whilst still remaining within the rigid stage gates required for tracking. Custom Dashboards were created bringing together information and links that were relevant to specific stages of the process. These gave tailored views to users for each part of the process providing information such as project ID, status, date created, linking the ELN pages associated with those projects. Dashboards were also used to obtain overall platform status metrics were used for tracking performance criteria and to generate kaizen improvement project ideas.

While SCD is a manual process it is used due to the simplicity, speed and applicability to most cell types. While fluorescence activated cell sorting (FACS) can be used to sort single cells into plates that can increase contamination, result in poor cell viability, is time consuming and relies on forward and side scatter which can be inaccurate in detecting cells.^18^ FACS also requires specialised trained staff and equipment to execute. Dedicated single cell dispensing instrumentation such as VIPS® (Advanced Instruments) and UP.SIGHT (BICO) can also be used produce clonal wells. Every well in a plate can obtain a single cell rather than a following the Poisson distribution associated with limiting dilution. The result is that fewer plates are required to obtain the desired numbers of clones. However, dedicated instruments can be unsuitable for operating a platform with large heterogenous types of cell line due to set up time and optimisations required for each cell type limiting speed and flexibility while requiring highly trained staff to optimise.

A key part of CLE was validating the clonality of engineered cells. Our platform utilised SCD and to support this an automated AI imaging platform was built to determine clonality which allowed for the capture of images from up to 240 384-well cell culture plates per day, or 96,160 images per day from a single instrument. For this number of generated images, manual and segmentation-based analysis methods were impractical for large numbers of brightfield images. Therefore, it was important to develop a software solution for image analysis that kept pace with the image generation rate of the automated system to prevent workflow bottlenecks. We developed an automated integrated imaging workcell and workflow data analysis pipeline in the cloud, together with a web application to manage the system. The platform produces a list of putative clones identified over the scanning period that are used to pick samples to move to the next stage.

The image capture workcell used GBG as the automation scheduling software. Other providers could include plate::works (Revvity) and Cellario (HiRes BioSolutions) which both have equal scheduling capabilities and drivers for the required instrumentation. GBG was chosen due to the flexibility in adding our custom .dll libraries that enabled our system to place messages on the Azure service bus that triggered worklfows to upload data and process it in the cloud. GBG was also chosen due to the flexibility to use equipment that we already had that was underutilised. Repurposing and reconfiguring equipment as needs change is becoming more common with companies offering self-deployed systems (HiRes), open-source schedulers (Galago) and drivers (UniteLabs).

Cell Metric® imager was chosen as the brightfield microscope to capture images of 384-well plates from SCD. The Cell Metric® was chosen due to the ability to capture whole well .TIFF images of entire 384 well plates in 3 mins without vignetting. The Celligo™ (Revvity) is equally fast and able to capture whole well images and can be integrated with the COBRA system described here. The Cytation 5 (Agilent) proved slower and prone to vignetting edge effects in plates.

Our approach was to use supervised ML to create a model so that the large amounts of data typically required for unsupervised learning was not needed. The Ilastik user interface used expert knowledge to mark-up areas of the images that contained cells and those that did not. Doing so gave the user instant feedback on how changes updated the classifications and made building an accurate model easier. A representative set of images was created for inconsistencies in the wells such as dust and marks resulting from the effects of irradiation that would occur over time. Other software systems such as CellProfiler and ImageJ could also have been used to produce classified brightfield images but those software lack the instant feedback of Ilastik^21^. Commercial software products such Phenologic.AI™ (Revvity) have also demonstrated the power of brightfield analysis in applications where cells can’t be stained, or in freeing up fluorophores to stain other cellular components.

We have, to date, autonomously collected and analysed over 16.8 million images. The model cannot identify a single cell in brightfield, but measures discrete colony counts down to 1% confluence. If images are captured at a high frequency at early stages, discreet small colonies can be resolved, and wells excluded if more than one is detected. Rather than producing a ‘black box’ model, our software designed to automate as much as possible any decision making, while retaining the ability to interrogate, interact with and QC samples to improve the quality of the desired outputs a feature which is essential for building trust and usability with lab based scientists.

Here we demonstrated how the interaction of configured software systems and custom programs that can represent physical process and data analysis, are equally as important tools as robotic automation for the development of HT CLE platforms. LIMS systems and robust data models facilitating a single source of truth allow for many useful paradigms or metrics to be created as questions arise. The platforms described have allowed for us to seamlessly manage multiple hundreds of projects per year up to 30 users simultaneously. The work demonstrates that when embarking on any large-scale automation project, building sufficient software infrastructure needs to be considered with similar attention as hardware. Without such systems, increased scale cannot be realised.

## Author Contributions

**David W. McClymont** contributed to conceptualization, data curation, funding acquisition, investigation, methodology, project administration, resources, software, supervision, validation, Writing - original draft and writing - review & editing. **Baird McIlwraith** contributed to investigation, software and writing - review & editing. **Sam Coulson** contributed to software and writing - review & editing. **Althea Green** contributed to investigation, methodology, project administration, validation, and writing - review & editing. **Elizabeth Scott** contributed to data curation, investigation, methodology and writing - review & editing.

## Data Availability Statement

No relevant data has been made available in this article.

